# The Bayesian Superorganism: Collective Probability Estimation in Swarm Systems

**DOI:** 10.1101/468942

**Authors:** Edmund R. Hunt, Nigel R. Franks, Roland J. Baddeley

## Abstract

Superorganisms such as social insect colonies are very successful relative to their non-social counterparts. Powerful emergent information processing capabilities would seem to contribute to the abundance of such ‘swarm’ systems, as they effectively explore and exploit their environment collectively. We develop a Bayesian model of collective information processing in a decision-making task: choosing a nest site (a ‘multi-armed bandit’ problem). House-hunting *Temnothorax* ants are adept at discovering and choosing the best available nest site for their colony: we propose that this is possible via rapid, decentralized estimation of the probability that each choice is best. Viewed this way, their behavioral algorithm can be understood as a statistical method that anticipates recent advances in mathematics. Our nest finding model incorporates insights from approximate Bayesian computation as a model of colony-level behavior; and particle filtering as a model of *Temnothorax* ‘tandem running’. Our framework suggests that the mechanisms of complex collective behavior can sometimes be explained as a spatial enactment of Bayesian inference. It facilitates the generation of quantitative hypotheses regarding individual and collective movement behaviors when collective decisions must be made. It also points to the potential for bioinspired statistical techniques. Finally, it suggests simple mechanisms for collective decision-making in engineered systems, such as robot swarms.

## Introduction

Of fundamental importance to most animals is finding food, mating opportunities, and obtaining a good place to live. The nests of social insects can confer considerable advantages to the colony’s defence, shelter, microclimate, and organization of function. While social insects like termites or paper wasps build their nest from scratch, certain house-hunting ants search the environment for pre-existing cavities that can be adapted to their needs. This entails the evolution of an effective collective exploration and decision-making procedure that nevertheless relies on the following of simple rules by individuals making use of local information. At the same time, in recent years Bayesian methods are increasingly recognised not only as useful *tools* for building descriptive models of biological phenomena, but that living systems *themselves* can be seen as Bayesian inference machines. As such, particular mechanisms in the exploration and decision-making process could correspond to Bayesian calculation of probabilities.

Two collective house-hunting systems have been investigated in detail: honeybee swarms looking for a new hive (Seeley and Buhrman, 2001) and *Temnothorax* ants (Franks et al., 2003b). In the case of *Temnothorax albipennis* ants, even when the nest is intact, colonies still send out scouts looking for better alternatives with an intensity inversely proportional to the quality of their current site (Doran et al., 2013), moving the whole colony if an improvement is identified (Dornhaus et al., 2004). Scouting ants identify candidate nests and recruit other ants one-by-one to inspect it by a process known as tandem running (Franks et al., 2006); when enough ants are at a site, a quorum is reached and emigration begins (Pratt, 2005; Pratt et al., 2002). Ants have clearly revealed preferences for what they like in a nest (Franks et al., 2006, 2003b), and artificial nest choice experiments can be easily conducted in the laboratory. Colonies are very good at discerning even small differences in important attributes such as nest darkness (Sasaki et al., 2013). Finding nest sites is a difficult challenge, and having to select between multiple options at the group level compounds the problem. Nevertheless, *T. albipennis* is an example of an ant species that does this very effectively: for example, they are able to choose a distant superior nest over an in-the-way poor one (Franks et al., 2008). The collective decision-making challenge of the ant colony parallels the same challenge for robot swarm systems, and as such a model of the biological process can be instructive for engineered swarm systems.

## Bayes and the Superorganism

The Bayesian perspective on animal behavior models individuals as having prior beliefs about the state of the world, expressed by probabilities, that are updated according to Bayes’ rule as new observations are made. Recent work on collective animal behavior uses this approach to weight non-social information (sensory experience, memory, internal states etc.) with social information (observations of others’ behaviors) in estimating probabilities; and a probability matching rule for making choices (Pérez-Escudero and de Polavieja, 2011; Arganda et al., 2012; Pérez-Escudero et al., 2013). Such work naturally has a focus on the individual, including for fish and humans (Pérez-Escudero and De Polavieja, 2017; Eguíluz et al., 2015; Mann, 2018), because individuals ‘selfishly’ optimise for their own advantage; this contrasts with highly related ant colonies, for example, which can properly be described as a form of ‘super-organism’ (Hölldobler and Wilson, 2009). In the superorganism, one can expect sophisticated collective strategies to emerge that are optimised for group-level performance. This distinguishes our approach: a focus on group-level, emergent Bayesian behavior (Models 2 and 3 below), in comparison to past work concerned with the individual Bayesian animal.

## The Multi-Armed Bandit

The house-hunting decision problem can be fruitfully mapped to the mathematical framework known as the ‘multi-armed bandit’ problem (MAB), whereby a player (or social unit: the ant colony) faces a choice of projects (nest locations) of unknown value. The name derives from the nickname ‘one-armed bandit’ for slot machines in a casino, where initially one does not know what the probability of making a payoff is for each machine but can only estimate this by repeated play. The MAB has been used to examine the foraging behavior of animals such as birds (Sherratt, 2011), bees (Keasar et al., 2002), fish (Thomas et al., 1985), and slime moulds (Reid et al., 2016). Making choices sequentially, the objective is to maximise the overall reward, which necessitates a trade-off between exploration (observing new locations to ascertain their quality), and exploitation (staying at the current location and making use of its benefits, but missing out on the potential gains to be had elsewhere). This trade-off is ubiquitous across biological domains, and is a key evolutionary force in the development of cognition (Hills et al., 2015).

The multi-armed bandit (MAB) is difficult to analyse as a theoretical problem to find the optimum solution, though in certain special cases this can be computed (Gittins, 1979). A ‘Gittins index’ may be calculated and then the problem simply becomes choosing the option with the highest index value (Gittins et al., 2011).Yet there remains a possibility of settling on playing the wrong arm forever, known as incomplete learning (Brezzi and Lai, 2000). This is analogous to the ants settling upon a good quality nest, when an excellent one is available nearby lying undiscovered.

### Thompson sampling

Optimal solutions to the MAB are difficult to compute, especially when there are many available options. One decision-making rule of thumb is known as Thompson sampling (Thompson, 1933), also called randomised probability matching (Scott, 2010). It has been rediscovered several times independently in the context of reinforcement learning (Ortega and Braun, 2010; Strens, 2000). Thompson sampling draws on Bayesian ideas, whereby the player (colony) has an assumed set of prior payoff distributions for each arm, and after making observations updates these to a posterior distribution using Bayes’ rule. The use of Bayes’ theorem and its assumption of prior ‘opinions’ about outcomes has been argued to be reasonable in the context of animal behavior, given both animals’ past individual experience of the environment, and the adaptation to that environment by previous generations (McNamara et al., 2006). In Thompson sampling, the player makes a random draw from all of these distributions, and chooses the arm (location) associated with the largest of these draws for its next observation. By making random draws from the player’s own distributions, which become more accurate with further observations, this decision-making rule permits the player mostly to choose the arm it believes to be the best. It also leaves open the possibility of making further observations from what the player believes to be a lower quality arm, if by chance it throws up a high sample value. In the case of searching for a new nest site, scouting ants are looking for the global quality maximum, and thus do not want to waste time returning to locations they know to be of lesser quality relative to what they have already observed. In a world of noisy observations, however, it can be imprudent to rule out an option prematurely. The Thompson sampling strategy achieves an effective compromise for this problem, and minimises what is known as ‘regret’ – the performance relative to a theoretical optimum (Agrawal and Goyal, 2012). This effectiveness is borne out in empirical studies of its effectiveness (Chapelle and Li, 2011), and it has been claimed to be a good model of humans responding to the ‘restless’ MAB problem, when the payoffs change through time (Speekenbrink and Konstantinidis, 2015). Thus, we hypothesize that *individual* ant behavior should approach a strategy resembling Thompson sampling (which is akin to probability matching).

### Approximate Bayesian computation

Approximate Bayesian computation (ABC) methods seek to bypass the evaluation of the likelihood function in Bayes’ rule in determining the posterior probability distribution. All ABC-based methods approximate the likelihood function by simulations, the outcomes of which are compared with observed data. The ABC rejection algorithm samples a set of parameter points from the prior distribution, simulates data sets under the statistical model specified by each of those parameter samples, and rejects those samples whose simulated data set is too different from the observed data set (summarised by a measure such as its mean), based on some distance measure between them (Turner and Van Zandt, 2012). This gives an approximated posterior distribution of accepted samples. In the case of ants, they make an observation of a location, and ‘reject’ it if it is not sufficiently high quality and move elsewhere, or accept it (stay there) if it is high quality. Individual *T. albipennis* ants do not need to make direct comparisons between alternative choices for the colony to choose collectively the best nest site (Robinson et al., 2009). An ant’s current location (which is accepted or rejected) is like one of the sampled parameter values in the described ABC rejection algorithm, and the judgement of sufficiently high quality is like the evaluation of the simulated data set against some tolerance level. The resulting ant locations (parameter sample) are an approximation of the required posterior probability – in the ants’ case, the probability that a location is the best, taking into account the ants’ observations. Hence, we argue that ABC is a good model of the *colony-level* strategy to solving the MAB: a decentralized computation of the probability that each option is the best one.

Recent work (Sasaki et al., 2018) finds that *T. rugatulus* colony decision-making behavior to be consistent with the *Sequential Choice* model, where competing options are evaluated in parallel, in a race to a decision threshold; whereas single ants match the *Tug of War* model where alternatives are directly compared and take longer as a result. This paper proposes an explanation for how this emergent change in behavior is possible: how individual rejection sampling can lead to a computationally meaningful macroscopic distribution of ants.

We develop a sequence of models in turn, before presenting results from simulations of those models and discussing their features. These include (1) effective individual sampling; (2) many such individuals sampling in parallel; (3) occasional communication between individuals. These models do not include a model of movement or pheromone communication. However, see Baddeley et al. 2019 for our Markov chain Monte Carlo movement model and Hunt et al. 2020 on the value of pheromone coordination (stigmergy): the present paper and these two other parts can be combined for a complete framework.

## Models and simulations

We develop three models that include progressively more behavioral features of an ant colony and become more effective in quickly identifying the highest quality potential nest location. Model 1 has (to motivate the problem) one ant trying to find the highest quality potential nest site out of three available. Model 2 then introduces the concept of approximate Bayesian computation (ABC) to explain the action of multiple house-hunting ants, and this model is found to identify the highest quality site more quickly than in the single ant case. Finally, in Model 3 tandem running is introduced whereby multiple ants explore in parallel, independently; but if a set quality threshold is exceeded in an ant’s estimation, another ant (randomly chosen) also observes that location at the next time iteration. Along with the ABC character of the ants’ decision-making process, this allows the highest quality location to be rapidly identified. We present individual runs of Models 2 and 3, and the average of 1000 simulations.

### Model 1: Single ant, Thompson sampling

Although our focus is on the group level, we begin with a simple model of the ant nest site selection process. There is a single ant and three potential locations for the ant to locate the nest. The existence of these three locations is known *a priori*, and there are no locations unknown to the ant. Although this model uses a form of reinforcement learning, a simple sequential accept/reject according to a fixed threshold would also be sufficient to proceed to Model 2.

The ant *i* (*i* = 1 only in this model) begins with a set of identical prior estimates of the quality of each location *L*. These are expressed as a separate mean and standard deviation for a normal distribution, *µ*_*iL*_ and *σ*_*iL*_, for each location *L* = 1, 2, 3. The uncertainty *σ*_*iL*_ expresses how confident the ant is in its estimate, and reduces with repeated observations. We assume that the ants start at a location that is not one of the three options: for example, their nest has been destroyed. However, a known starting location could be included with correspondingly low *σ*_*iL*_ (i.e. high certainty in its quality). The available locations have a true and constant quality *Q*_*L*_ that is accessible to the ant by making repeated noisy observations. These observations are drawn from a normal distribution with a mean *Q*_*L*_ and a standard deviation which is the observation error, *σ*_*obs*_, a fixed parameter ‘known’ to the ant. This encapsulates the fact that the methods ants use to find and assess the characteristics of a potential nest are associated with noise and error.

The ant makes a sequence of observations of the different locations to try and find the one with the highest quality. To decide which location to travel to and observe next, it draws a sample simultaneously from each of its subjective location quality estimates, 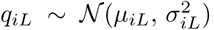, and moves to the location with the highest drawn sample. This is a form of Thompson sampling, or randomised probability matching. While this will tend to move the ant to the location with the highest quality estimate *µ*_*iL*_, its probabilistic nature means that sometimes a location with a lower quality estimate will generate a sample higher than that from the better-quality estimates, especially if the uncertainty in that estimate *σ*_*iL*_ is still relatively high. When the ant makes an observation, for instance of Location 3, it updates its estimate of the location’s quality using Bayes’ rule. With a known observation error *σ*_*obs*_, after making an observation of Location 3, *O*_*i*3_, the posterior distribution of nest quality *q*_*i*3_ for ant *i* can be updated according to a Kalman filter, as:

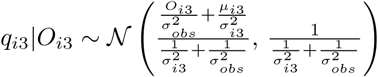

This is a weighted average of the quality value *O*_*i*3_ observed by ant *i*, with the ant’s prior estimate of it *µ*_*i*3_, according to the associated observation error *σ*_*obs*_, and the estimate of the error in *µ*_*i*3_ given by *σ*_*i*3_. If for instance the measurement error is low and the prior uncertainty was high, the posterior estimate of *µ*_*i*3_ will be close to the observed quality value. The above procedure of making a biased exploration based on current quality estimates, and updating those estimates based on observations of consequently chosen locations, can be iterated through a series of observations and will converge through time upon the true location quality *Q*_3_. Many animals exhibit behavior consistent with such Bayesian updating of environmental parameter estimates (Valone, 2006).

The ant’s running estimates of *µ*_*iL*_ and *σ*_*iL*_ for each location *L* = 1, 2, 3 encodes a summary statistic that captures all that an ant knows about each of the qualities at each step in time, so no memory of the individual past observations is needed. As more observations are made, the uncertainty *σ*_*iL*_ in the estimate of each location’s quality reduces toward zero, and thus it becomes less and less likely that the sampling step will lead the ant to revisit locations of lower quality. Eventually the ant stays in a particular location, provided it has a quality higher than the other alternatives, and this represents a choice being made in favour of that location.

Although the ant in this model can make its choice fairly quickly, because there are only three locations to visit, it still requires several observations to be made. In practice, house-hunting ants like *T. albipennis* make their choice using several scouting ants, after each makes only a few observations of one or two of the potential locations: and it can still make accurate choices because it makes its decisions collectively with a quorum sensing mechanism (Pratt et al., 2002). The next model shows how this is analogous to approximate Bayesian computation (ABC).

### Model 2: Multiple ants, ABC

In this model, individual ants make observations in the same way as in the single ant model, and do so independently. However, their global distribution across the three locations is used as a running estimate of the posterior (post observation) probability that each location is the best available place to establish the colony’s nest. We argue that this is analogous to approximate Bayesian computation (ABC).

As an example, we can simulate five exploring ants, which for a period of time make observations and decide at each time step whether to ‘accept’ a site (stay in the location, exploitation) or ‘reject’ a site (move to a new location, exploration). Note, however, that rejection may be temporary: the ant may reconsider and return to previously visited locations; though with better alternatives available a return to a low-quality location becomes increasingly improbable. At the end of the time period there are four ants at Location 3, one ant at Location 2, and no ants at Location 1 (see Fig. 1).

**Figure 1:**
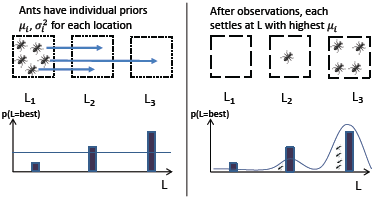
Multiple non-interacting ants engage in Thompson sampling until they settle on the location each believes is the highest quality. Their macroscopic distribution approximates the true posterior distribution of location qualities.

#### Algorithm 1: Pseudocode for simple approximate Bayesian computation nest finding method.

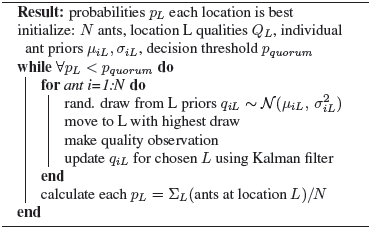

This corresponds to a probability of 80% (four out of five ants) that Location 3 is the best location; similarly 20% and 0% for locations 2 and 1. Psuedocode for this algorithm is provided in Algorithm 1.

The ants employ a quorum sensing mechanism (Pratt et al., 2002), whereby a decision is taken in favour of a location when individual ants have a high enough encounter rate with other ants, indirectly estimating the number of ants at that location (Pratt, 2005). The quorum threshold is introduced simply by specifying a probability *p*_*quorum*_, which is exceeded when enough ants are at a location (excluding the known starting location of a destroyed nest). Here we consider an example threshold of *p*_*quorum*_ = 0.7. When the need to identify a new nest is urgent, for instance when the colony’s nest is damaged, a high threshold will be a handicap. In the case where a new nest is needed quickly, the quorum threshold for a decision is lowered, sometimes at the expense of accuracy (in terms of choosing the best option) (Franks et al., 2003a); this speed-accuracy trade-off is faced by animals in many contexts (Chittka et al., 2009). As a result, the medium quality site may be estimated as the best site. Nevertheless, the Thompson–ABC model presented here would still be a good representation of this aspect of the ants’ behavior.

The global distribution of a sufficient number of ants (ten, for example) is an approximation of the posterior probability that each choice (potential nest site, for instance) is the best available; when a threshold probability (resource quality) is exceeded a decision can be said to have been reached. Allowing many ants to explore in parallel allows a decision to be taken more quickly, because while the total amount of ant exploration time (number of ants *×* time) may be similar, the actual time taken can be much less. This is important for biological feasibility, since an individual ant cannot explore indefinitely, and is unlikely to discover all available sites independently. This ABC model captures the emergent, statistical phenomena whereby the spatial distribution of scouting ants determines the collective decision. However, one obvious simplification made is that the ants are non-interacting, such that each ant has to engage in a costly search process (in terms of time and energy) and make its own observations of a location that may have already been visited many times by its nestmates. Pheromone marking can facilitate coordination (Franks et al., 2007; Sasaki et al., 2014; Hunt, 2020), as can tandem running.

### Model 3: ABC with particle filtering

Model 3 adds a representation of the *T. albipennis* tandem running behavior, whereby a scouting ant that has found a high-quality resource (like a potential new nest site) finds another worker ant and attempts to lead her to that spatial position. This is illustrated in Fig. 2. Interaction between individuals can contribute significantly to an emergent collective sensing of the environment that does not require enhancement of individual abilities (Berdahl et al., 2013). The tandem running behavior is implemented in the model by assigning each scouting ant a threshold quality, whereby if their estimate of their current location’s quality exceeds that threshold, they are triggered to attempt the leading of another scouting ant to that location. The tandem run behavior can be understood as a form of ‘particle filter’ method (also known as Sequential Monte Carlo) (Speekenbrink, 2016). Such methods boost sampling from higher quality (probability) space. The threshold *q*_*lead*_ is identical for all ants in this model (though see Discussion). When an ant observes a high-quality location, resulting in an estimate that exceeds *q*_*lead*_, another ant from all of those exploring is randomly chosen for an attempted recruitment. If that ant’s estimate of its current location’s quality is lower than or equal to the leader ant’s estimate of the proposed location, it relocates to the leader ant’s position and makes a new observation of it. If the quality of the tandem leader’s location, as estimated by the leader, is less than that estimated by the potential follower at its current position, it resists recruitment and stays put. Pairwise comparison between two options has been referred to as a ‘duelling bandits’ scenario (Yue et al., 2012), although in this case there are two agents (proposer and follower) rather than one observer.

**Figure 2:**
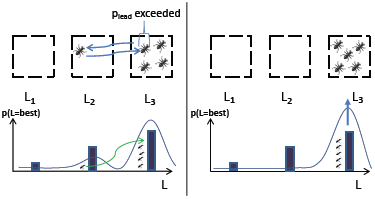
The tandem running behavior acts like a particle filter, boosting sampling from higher quality locations. Ants, having discovered a high-quality location, recruit nestmates to also observe that location. This results in a posterior distribution (global distribution of ants) that is more likely to converge upon the true state of the environment (right pane).

## Results

### Model 1: Single ant, Thompson sampling

An example simulation of Model 1 is shown in Fig. 3. The ant starts with three identical priors over the three locations, *µ*_*i*=1:3_ = 1, *σ*_*i*=1:3_ = 0.5, makes observations with fixed, known error *σ*_*obs*_ = 0.5, while the actual location qualities are set as *q*_1_ = 1, *q*_2_ = 1.1, *q*_3_ = 1.2. As per the Thompson sampling procedure, it draws samples from its estimated quality distributions, and moves to the location producing the highest sample, where it makes an observation that decreases its uncertainty in the estimate. Over the course of 100 time steps in this example, its location changes several times, until it narrows its search to locations 1 and 2 from *t* = 14 onwards. It then settles on Location 3 as the highest quality from *t* = 56, when it does not move any more (the samples from its estimated quality distributions always produce Location 3 as the highest quality). Its running estimate of the location qualities is shown in the second pane in Fig. 3; its estimate for locations 2 and 3 becomes accurate as it makes repeated observations, while its lack of observations for Location 3 means its quality is somewhat underestimated. The uncertainty in the ant’s estimates for locations 2 and 3 steadily decreases as new observations are made. Because the ant acts alone it must make several observations of each location before it can be confident that it has identified one with the highest available quality. In practice, an advantage of the colony’s tight-knit social organization is that multiple ants can explore and observe different locations in parallel, greatly accelerating the process.

**Figure 3:**
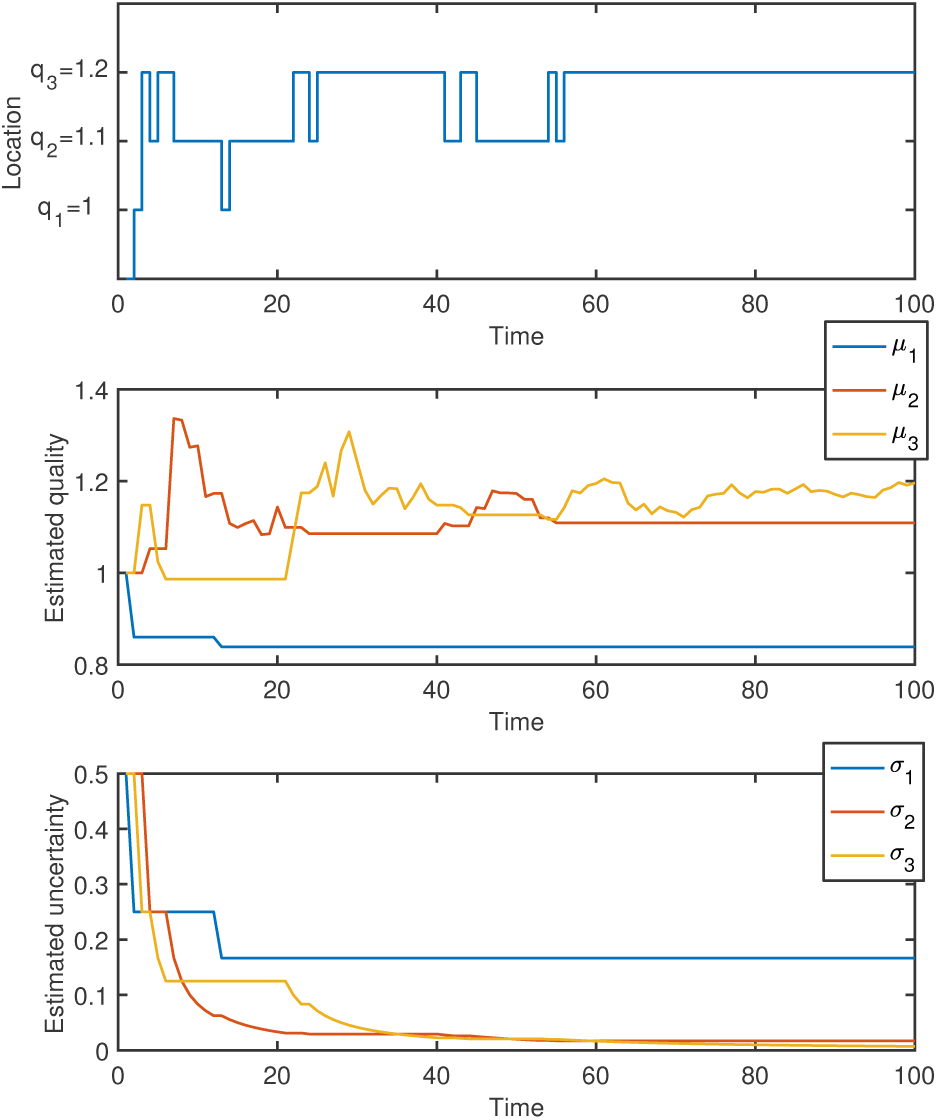
Model 1: Single ant using Thompson sampling to identify the best nest. Location 3 has the highest quality, and after sampling the locations the ant makes several observations from it and accurately estimates its quality, *q*_3_ = 1.2, and stays in that location after around 60 iterations.

### Model 2: Multiple ants, ABC

With the Thompson sampling procedure working as before, this model initiates multiple ants exploring in parallel. Fig. 4 shows the result of simulating ten ants for 100 time steps, with the number of ants in each location shown on the y-axis. This number corresponds to the probability that each location is the best, when normalised, from the level of the colony’s ‘collective cognition’. In this case, a threshold of 60% probability (6 of 10 ants) is breached quickly at *t* = 4 for Location 2, and then 80% at Location 3 at *t* = 6. This is followed by a clear lead for Location 3 at around *t* = 30. At the end of 100 simulated time steps, the model has settled on probabilities of 80%, 20% and 0% for each of locations 3, 2 and 1 being the best available quality.

**Figure 4:**
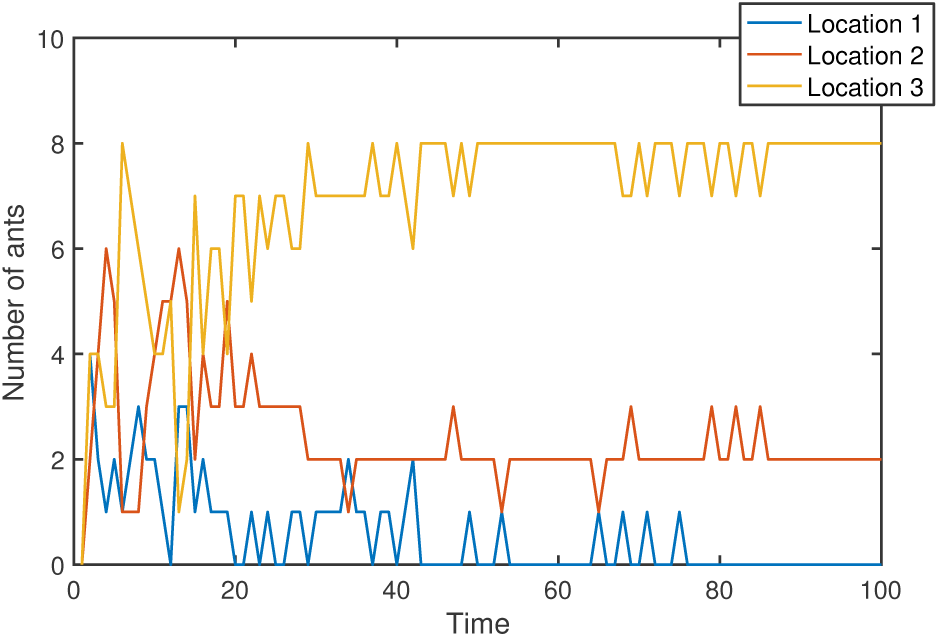
Model 2: Multiple (ten) ants independently exploring three locations. Their global distribution approximates the probability that each location is the highest quality available. At t=100, there are 8, 2, and 0 at locations 3, 2, and 1, or probabilities of 80%, 20%, and 0% that these are the best choices. A clear lead for Location 3 is established much earlier at around *t* = 30.

This model highlights two aspects of real ant colony decision-making: first, that working collectively allows the optimal choice to be identified more quickly; but second, that it is possible for premature decisions to be made in favour of a site of good quality (Location 2) over the site of the best quality (Location 3) if a threshold is set too low or breached too quickly. For example, with a threshold of 60% probability a quorum in favour of Location 2 may have been met at *t* = 4. It is important, then, to identify high quality locations quickly, otherwise they may be discovered too late in the nest choice process; and to set a suitable quorum threshold.

### Model 3: ABC with particle filtering

The final model includes a representation of the *T. albipennis* tandem running behavior to preferentially sample from high quality regions. An example of the model’s output, for the same location qualities as before, is shown in Fig. 6. A high estimated quality threshold of *q*_*lead*_ = 1.25 is set in this case. The potential follower will move to the high-quality location only if the leader’s estimate of that location’s quality is higher than the follower’s estimate of her current position’s quality. After some initial volatility, as ants switch between location 2 and 3, a clearer preference for location 3 is established than in Model 2 (c.f. Fig. 4) after around *t* = 25. Comparing the average performance of Models 2 and 3 (Fig. 5 and 7), a quorum of 70% is reached much more quickly: *t* = 34 with the tandem running vs. *t* = 91 without, less than half the time.

**Figure 5:**
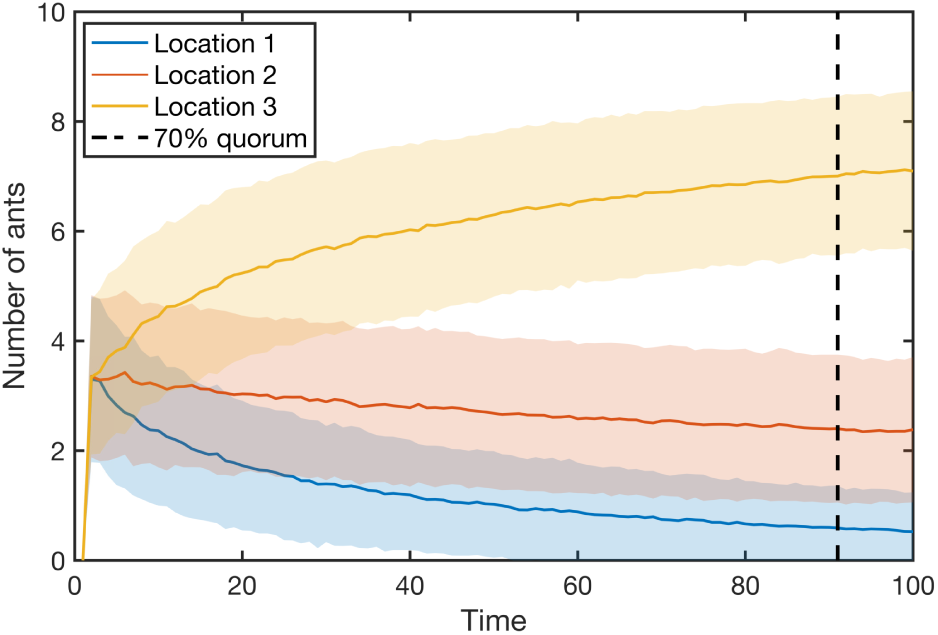
Model 2: Average number of ants at each location over 1000 simulations. A 70% quorum is reached at *t* = 91 on average. At *t* = 100, there are 0.52, 2.38 and 7.09 ants at Locations 1-3 respectively. Bands are standard deviation.

**Figure 6:**
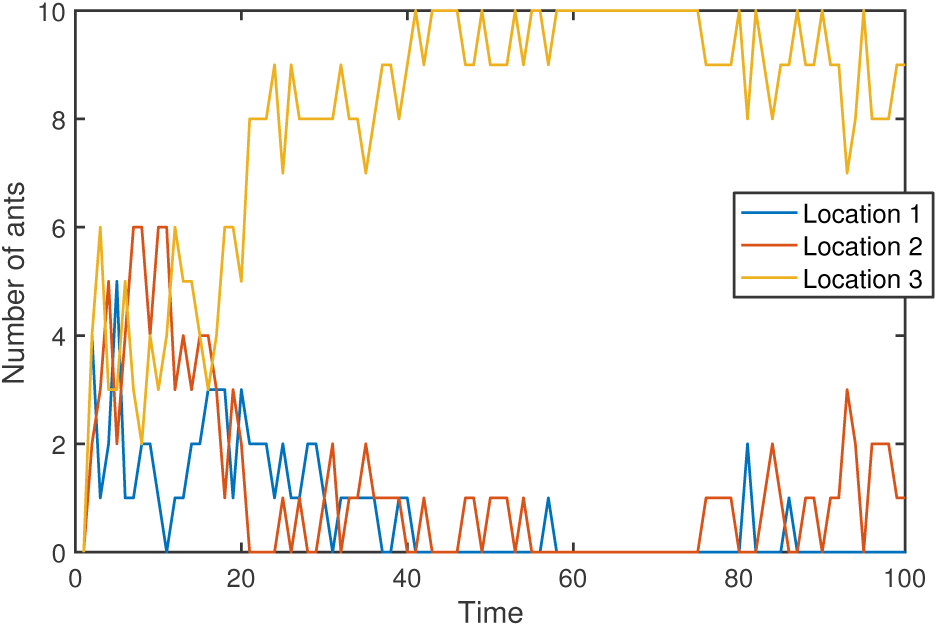
Model 3: Ten exploring ants, with a tandem running (‘particle filter’) behavior included. Here, more ants reach Location 3, more quickly.

**Figure 7:**
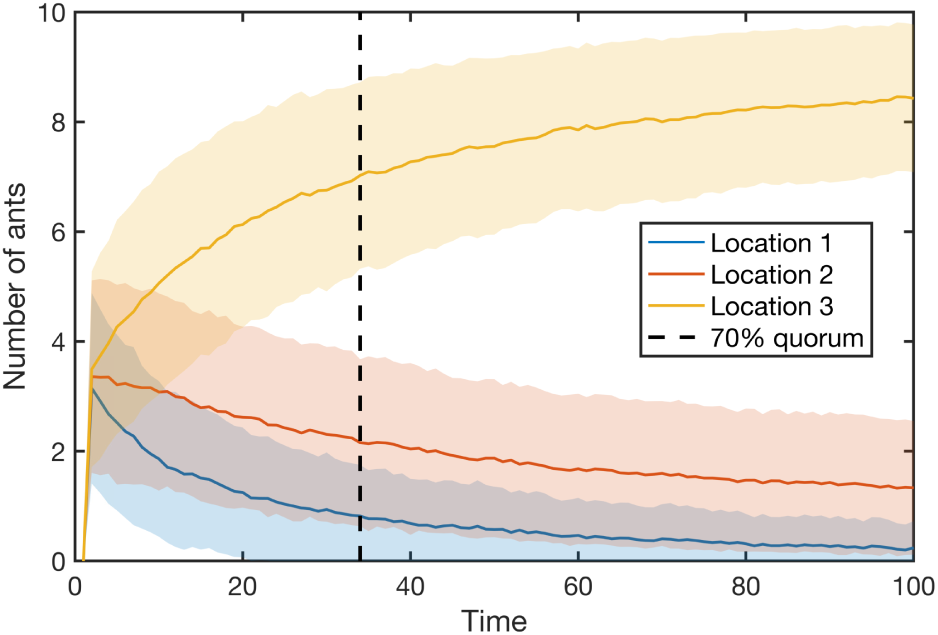
Model 3: Average number of ants at each location over 1000 simulations. A 70% quorum is reached at *t* = 34 on average. At *t* = 100, there are 0.24, 1.33 and 8.43 ants at Locations 1-3 respectively. Bands are standard deviation.

## Discussion

Three models of ‘swarm’ decision-making (*Temnothorax* ant colony nest choice) have been developed that progressively include more realistic features, and that are more efficient and effective at selecting the best available nest site location. This ‘multi-armed bandit’ problem is solved through individual-level Thompson sampling, and colony-level probability estimation that we argue is analogous to approximate Bayesian computation. Tandem running (‘particle filtering’) is introduced in the final model with multiple ants to ensure that information about the position of high quality locations is shared among the ants. This necessitates the introduction of a new parameter *q*_*lead*_, which is a threshold quality above which tandem runs are initiated by the observer. While an appropriate *q*_*lead*_ and *p*_*quorum*_ are set in this model by reference to the known overall distribution of qualities, in nature the ants do not know the distribution of location qualities *a priori*. However, the ants’ evolutionary ‘experience’ should favour the setting of thresholds that are appropriate to their typical environment. This is because colonies that have made beneficial use of e.g. the tandem running behavior to help them reproduce are likely to have avoided both the pitfalls of *q*_*lead*_ being too high (so the behavior is never used) and *q*_*lead*_ being too low (excessive sharing of low value information). Phenotypic plasticity should also adapt colonies to actual environmental conditions (Hunt, 2020). A heterogeneous distribution of *q*_*lead*_ is likely, and perhaps even desirable, since it allows information of variable value to be propagated through the colony and collectively assessed. The optimal and empirical distribution of thresholds in social insects is an active topic of research (e.g. (Robinson et al., 2012)) and models have been developed showing the effectiveness of heterogeneous thresholds (Masuda et al., 2015). A recent study of *T. rugatlus* identified four behavioral castes in the house hunting process, with a small proportion (3-12%) of the colony being ‘wandering workers’ (Valentini et al., 2020). These ants are inactive in the decision-making process (they do not recruit through tandem runs) but still are active in visiting nest sites. This would correspond to ants with a high *q*_*lead*_ in our model, and may help the colony to take advantage of a ‘move to improve’ opportunity if their ‘pickiness’ means an excellent quality nest is eventually found (Valentini et al., 2020; Dornhaus et al., 2004).

Our model suggests how collective cognition in the ‘superorganism’ can emerge via collective probability estimation, realised through a form of spatial approximate Bayesian computation. Future work can probe in more detail how far this analogy operates. Such models of superorganism decision-making behaviors may be successfully transposed into an engineering context, for example in relation to robot ‘swarms’ that are also inspired by social insect self-organization (Şahin, 2005).

## Acknowledgements

E.R.H. thanks the UK Engineering and Physical Sciences Research Council (grants no. EP/I013717/1 to the Bristol Centre for Complexity Sciences and EP/N509619/1, DTP 2016-17 University of Bristol, EPSRC DTP Doctoral Prize).

## References

Agrawal, S. and Goyal, N. (2012). Analysis of Thompson Sampling for the Multi-armed Bandit Problem. In 25th Annual Conference on Learning Theory, pages 39.1–39.26. Journal of Machine Learning Research.

Arganda, S., Pérez-Escudero, A., and de Polavieja, G. G. (2012). A common rule for decision making in animal collectives across species. Proceedings of the National Academy of Sciences, 109(50):20508–20513.

Baddeley, R. J., Franks, N. R., and Hunt, E. R. (2019). Optimal foraging and the information theory of gambling. Journal of The Royal Society Interface, 16(157):20190162.

Berdahl, A., Torney, C. J., Ioannou, C. C., Faria, J. J., and Couzin, D. (2013). Emergent Sensing of Complex Environments by Mobile Animal Groups. Science, 339(6119):574–576.

Brezzi, M. and Lai, T. L. (2000). Incomplete Learning from Endogenous Data in Dynamic Allocation. Econometrica, 68(6):1511–1516.

Chapelle, O. and Li, L. (2011). An empirical evaluation of thompson sampling. In Advances in Neural Information Processing Systems, pages 2249–2257.

Chittka, L., Skorupski, P., and Raine, N. E. (2009). Speed-accuracy tradeoffs in animal decision making. Trends in Ecology & Evolution, 24(7):400–407.

Doran, C., Pearce, T., Connor, A., Schlegel, T., Franklin, E., Sendova-Franks, A. B., and Franks, N. R. (2013). Economic investment by ant colonies in searches for better homes. Biology Letters, 9(5).

Dornhaus, A., Franks, N. R., Hawkins, R. M., and Shere, H. N. S. (2004). Ants move to improve: colonies of *Leptothorax albipennis* emigrate whenever they find a superior nest site. Animal Behaviour, 67:959–963.

Egúiluz, V. M., Masuda, N., and Fernández-Gracia, J. (2015). Bayesian decision making in human collectives with binary choices. PLoS ONE, 10(4):1–14.

Franks, N. R., Dornhaus, A., Best, C. S., and Jones, E. L. (2006). Decision making by small and large house-hunting ant colonies: one size fits all. Animal Behaviour, 72(3):611–616.

Franks, N. R., Dornhaus, A., Fitzsimmons, J. P., and Stevens, M. (2003a). Speed versus accuracy in collective decision making. Proceedings of the Royal Society B-Biological Sciences, 270(1532):2457–2463.

Franks, N. R., Hardcastle, K. A., Collins, S., Smith, F. D., Sullivan, K. M. E., Robinson, E. J. H., and Sendova-Franks, A. B. (2008). Can ant colonies choose a far-and-away better nest over an in-the-way poor one? Animal Behaviour, 76:323–334.

Franks, N. R., Hooper, J. W., Dornhaus, A., Aukett, P. J., Hayward, A. L., and Berghoff, S. M. (2007). Reconnaissance and latent learning in ants. Proceedings of the Royal Society of London B: Biological Sciences, 274(1617):1505–1509.

Franks, N. R., Mallon, E. B., Bray, H. E., Hamilton, M. J., and Mischler, T. C. (2003b). Strategies for choosing between alternatives with different attributes: exemplified by house-hunting ants. Animal Behaviour, 65(1):215–223.

Gittins, J., Glazebrook, K., and Weber, R. (2011). Multi-armed bandit allocation indices. John Wiley & Sons.

Gittins, J. C. (1979). Bandit processes and dynamic allocation indexes. Journal of the Royal Statistical Society Series B-Methodological, 41(2):148–177.

Hills, T. T., Todd, P. M., Lazer, D., Redish, A. D., Couzin, I. D., and Cognitive Search Res, G. (2015). Exploration versus exploitation in space, mind, and society. Trends in cognitive sciences, 19(1):46–54.

Hölldobler, B. and Wilson, E. O. (2009). The superorganism: the beauty, elegance, and strangeness of insect societies. WW Norton & Company.

Hunt, E. R. (2020). Phenotypic Plasticity Provides a Bioinspiration Framework for Minimal Field Swarm Robotics. Frontiers in Robotics and AI, 7:23.

Hunt, E. R., Franks, N. R., and Baddeley, R. J. (2020). The Bayesian Superorganism: externalised memories facilitate distributed sampling. Journal of the Royal Society Interface, (In press).

Keasar, T., Rashkovich, E., Cohen, D., and Shmida, A. (2002). Bees in two-armed bandit situations: foraging choices and possible decision mechanisms. Behavioral Ecology, 13(6):757–765.

Mann, R. P. (2018). Collective decision making by rational individuals. Proceedings of the National Academy of Sciences, 115(44):E10387–E10396.

Masuda, N., O’Shea-Wheller, T. A., Doran, C., and Franks, N. R. (2015). Computational model of collective nest selection by ants with heterogeneous acceptance thresholds. Royal Society Open Science, 2(6):140533.

McNamara, J. M., Green, R. F., and Olsson, O. (2006). Bayes’ theorem and its applications in animal behaviour. Oikos, 112(2):243–251.

Ortega, P. A. and Braun, D. A. (2010). A minimum relative entropy principle for learning and acting. Journal of Artificial Intelligence Research, 38:475–511.

Pérez-Escudero, A. and de Polavieja, G. G. (2011). Collective Animal Behavior from Bayesian Estimation and Probability Matching. PLoS Computational Biology, 7(11).

Pérez-Escudero, A. and De Polavieja, G. G. (2017). Adversity magnifies the importance of social information in decision-making. Journal of the Royal Society Interface, 14(136).

Pérez-Escudero, A., Miller, N., Hartnett, A. T., Garnier, S., Couzin, I. D., and de Polavieja, G. G. (2013). Estimation models describe well collective decisions among three options. Proceedings of the National Academy of Sciences, 110(37):E3466–E3467.

Pratt, S. C. (2005). Quorum sensing by encounter rates in the ant Temnothorax albipennis. Behavioral Ecology, 16(2):488–496.

Pratt, S. C., Mallon, E. B., Sumpter, D. J. T., and Franks, N. R. (2002). Quorum sensing, recruitment, and collective decision-making during colony emigration by the ant Leptothorax albipennis. Behavioral Ecology and Sociobiology, 52(2):117–127.

Reid, C. R., MacDonald, H., Mann, R. P., Marshall, J. A. R., Latty, T., and Garnier, S. (2016). Decision-making without a brain: how an amoeboid organism solves the two-armed bandit. Journal of the Royal Society, Interface / the Royal Society, 13(119).

Robinson, E. J. H., Feinerman, O., and Franks, N. R. (2012). Experience, corpulence and decision making in ant foraging. Journal of Experimental Biology, 215(15):2653–2659.

Robinson, E. J. H., Smith, F. D., Sullivan, K. M. E., and Franks, N. R. (2009). Do ants make direct comparisons? Proceedings of the Royal Society B-Biological Sciences, 276(1667):2635–2641.

Sasaki, T., Granovskiy, B., Mann, R. P., Sumpter, D. J. T., and Pratt, S. C. (2013). Ant colonies outperform individuals when a sensory discrimination task is difficult but not when it is easy. Proceedings of the National Academy of Sciences of the United States of America, 110(34):13769–13773.

Sasaki, T., Hölldobler, B., Millar, J. G., and Pratt, S. C. (2014). A context-dependent alarm signal in the ant Temnothorax rugatulus. Journal of Experimental Biology, 217(18):3229–3236.

Sasaki, T., Pratt, S. C., and Kacelnik, A. (2018). Parallel vs. comparative evaluation of alternative options by colonies and individuals of the ant *Temnothorax rugatulus*. Scientific Reports, 8(1):1–8.

Scott, S. L. (2010). A modern Bayesian look at the multi-armed bandit. Applied Stochastic Models in Business and Industry, 26(6):639–658.

Seeley, D. T. and Buhrman, C. S. (2001). Nest-site selection in honey bees: how well do swarms implement the “best-of-N” decision rule? Behavioral Ecology and Sociobiology, 49(5):416–427.

Sherratt, T. N. (2011). The optimal sampling strategy for unfamiliar prey. Evolution, 65(7):2014–2025.

Speekenbrink, M. (2016). A tutorial on particle filters. Journal of Mathematical Psychology, 73:140–152.

Speekenbrink, M. and Konstantinidis, E. (2015). Uncertainty and Exploration in a Restless Bandit Problem. Topics in Cognitive Science, 7(2):351–367.

Strens, M. (2000). A Bayesian framework for reinforcement learning. In International Conference on Machine Learning, pages 943–950.

Thomas, G., Kacelnik, A., and Van Der Meulen, J. (1985). The three-spined stickleback and the two-armed bandit. Behaviour, pages 227–240.

Thompson, W. R. (1933). On the likelihood that one unknown probability exceeds another in view of the evidence of two samples. Biometrika, 25:285–294.

Turner, B. M. and Van Zandt, T. (2012). A tutorial on approximate Bayesian computation. Journal of Mathematical Psychology, 56(2):69–85.

Valentini, G., Masuda, N., Shaffer, Z., Hanson, J. R., Sasaki, T., Walker, S. I., Pavlic, T. P., and Pratt, S. C. (2020). Division of labour promotes the spread of information in colony emigrations by the ant *Temnothorax rugatulus*. Proceedingsof the Royal Society B-Biological Sciences, 287(1924):20192950.

Valone, T. J. (2006). Are animals capable of Bayesian updating? An empirical review. Oikos, 112(2):252–259.

Yue, Y., Broder, J., Kleinberg, R., and Joachims, T. (2012). The K-armed dueling bandits problem. Journal of Computer and System Sciences, 78(5):1538–1556.

Şahin, E. (2005). Swarm Robotics: From Sources of Inspiration to Domains of Application. In Şahin, E. and Spears, W. M., editors, Swarm Robotics - SAB 2004 International Workshop. Lecture notes in computer science, pages 10–20. Springer, Berlin, Heidelberg.

